# TreeBASEdmp: A Toolkit for Phyloinformatic Research

**DOI:** 10.1101/399030

**Authors:** William H. Piel, Rutger A. Vos

**Affiliations:** Yale-NUS College, 10 College Avenue West #01-101, Singapore 138609; National University of Singapore, Department of Biological Sciences, Singapore.; Naturalis Biodiversity Center, P.O. Box 9517, 2300RA Leiden, the Netherlands; Leiden University, Institute of Biology Leiden, the Netherlands

**Keywords:** TreeBASE, phyloinformatics, supertree, SQL, phylogenetic querying

## Abstract

Over the last 20 years, TreeBASE has acquired a substantial body of phylogenetic data, including more than 20,000 published phylogenies. Given latency issues and limited options when it comes to querying the database remotely, a simplified and consolidated version of the database, here called TreeBASEdmp, is made available for download, allowing biologists to design custom analyses of the data on their local computers. The database is indexed to support searching for phylogenetic topologies using nested sets and closure tables. Here we propose a new approach to find broadly-defined phylogenetic patterns, a method we call *Generic Topological Querying*, which allows the user to find hypotheses of relationship without being constrained to use particular sets of specific taxa. Additionally, we normalize as many leaf nodes as possible to an equivalent species rank identifier to assist in supertree synthesis. Our example script rapidly assembles sets of trees and generates a matrix representation of them for subsequent supertree generation.

---

Growth in phylogenetic data is akin to Moore’s Law, being the product of faster computers, speedier DNA sequencers, and more efficient algorithms. The result is near-exponential growth in phylogenetic knowledge (Sanderson, M.J., Baldwin, B., et al%. 1993, Pagel, M. 1997, Pagel, M. 1999, Page, R.D.M. 2005, Parr, C.S., Guralnick, R., et al. 2012), with the promise that subsequent synthesis and meta-analysis of these data will bring forth a “new age of discovery,” among other benefits (Donoghue, M.J. and Alverson, W.S. 2000, Soltis, P.S. and Soltis, D.E. 2001, Cracraft, J. 2002, McTavish, E.J., Drew, B.T., et al. 2017). To attain this goal, standards for data serialization such as NeXML (Vos, R.A., Balhoff, J.P., et al. 2012), phyloXML (Han, M.V. and Zmasek, C.M. 2009), and NEXUS (Maddison, D.R., Swofford, D.L., et al. 1997) have been developed; ontologies such as CDAO (Prosdocimi, F., Chisham, B., et al. 2009) have been defined; minimum reporting standards such as MIAPA (Leebens-Mack, J., Vision, T., et al. 2006) have been proposed, and interfaces for phyloinformatic web services, such as the PhyloWS API, have been drafted (Lapp, H. and Vos, R.A. 2009). Software libraries that make use of these standards, such as Perl (Vos, R.A, Caravas, J., et al. 2011), Python (Sukumaran, J. and Holder, M.T. 2010, Talevich, E., Invergo, B.M., et al. 2012), and R (Boettiger, C. and Temple Lang, D. 2012) provide computational environments for automating large-scale phyloinformatic analysis. Collectively, these technologies create a cyberinfrastructure conducive to meta-analysis on the growing body of phylogenetic results.

In order to synthesize and analyze phylogenetic knowledge, none of these technologies would be particularly useful without first compiling the cumulative output of phylogenetic research in digital and reusable form. Some twenty years ago, TreeBASE was developed to archive phylogenetic data, else these data would be “lost” in the printed literature or erased from researchers’ hard drives (Sanderson, M., Donoghue, M., et al. 1994, Piel, W.H., Donoghue, M., et al. 2002). Since submission to the database is largely voluntary, TreeBASE has only managed to captured a fraction of the totality of phylogenetic effort (Page, R.D.M. 2005, Parr, C.S., Guralnick, R., et al. 2012), however even a relatively small sample of trees has been shown to provide fairly broad tree connectivity, suggesting that supertree assembly of the tree of life is achievable despite incomplete data (Sanderson, M.J., Purvis, A., et al. 1998, Piel, W.H. 2003, Piel, W.H., Sanderson, M.J., et al. 2003). Additionally, TreeBASE has provided a core dataset that researchers can use to investigate how to improve taxonomic (Herbert, K.G., Gehani, N.H., et al. 2004, Herbert, K.G., Pusapati, S., et al. 2005, Page, R.D.M. 2006, Page, R.D.M. 2007, Anwar, N. and Hunt, E. 2009, Ranwez, V., Clairon, N., et al. 2009) and topological querying or browsing (Shan, H., Herbert, K.G., et al. 2002, Wang, J.T., Shan, H., et al. 2003, Wang, J.T., Shan, H., et al. 2005, Chevenet, F., Brun, C., et al. 2006, Chen, D., Burleigh, J.G., et al. 2008, Hossain, S., Islam, M., et al. 2008, Chisham, B., Wright, B., et al. 2011, Le, T., Nguyen, H., et al. 2012). Finally, TreeBASE, supplemented with data from Dryad (White, H., Carrier, S., et al. 2008) served as the core source of data for Open Tree of Life, a platform for fusing phylogenetic trees with taxonomic classifications to produce a comprehensive tree of life (Hinchliff, C.E., Smith, S.A., et al. 2015).

Although TreeBASE is incomplete (Page, R.D.M. 2005, Parr, C.S., Guralnick, R., et al. 2012), this resource still represents a substantial quantity of data. Thus far about 5,000 registered submitters have uploaded data from about 6,000 publications and 600 different journals, crediting about 20,000 distinct author names. As of July 2018, these data include 15,223 matrices (with a total sum of 1,104,920 rows and 104,632,720 columns) and 20,246 trees (with a total of 1,311,445 leaf nodes). The leaf nodes map to a total of 117,231 distinct taxa in the NCBI taxonomy and 129,788 distinct uBio namebank records. The trees and matrices cover taxa in the following proportions: 34% Viridiplantae; 33% Fungi; 28% Metazoa; 3% Bacteria; 1% Archaea; and 1% Viruses. This taxonomic coverage is biased depending on the degree that different communities of biologists commit to submitting data. A comparison with the number of species in Genbank (Fig. 1) shows that when there are more species in Genbank, there are more trees in TreeBASE -- however some groups are overrepresented (e.g. Fungi) and some are underrepresented (e.g. prokaryotes and viruses).

**Figure 1.**
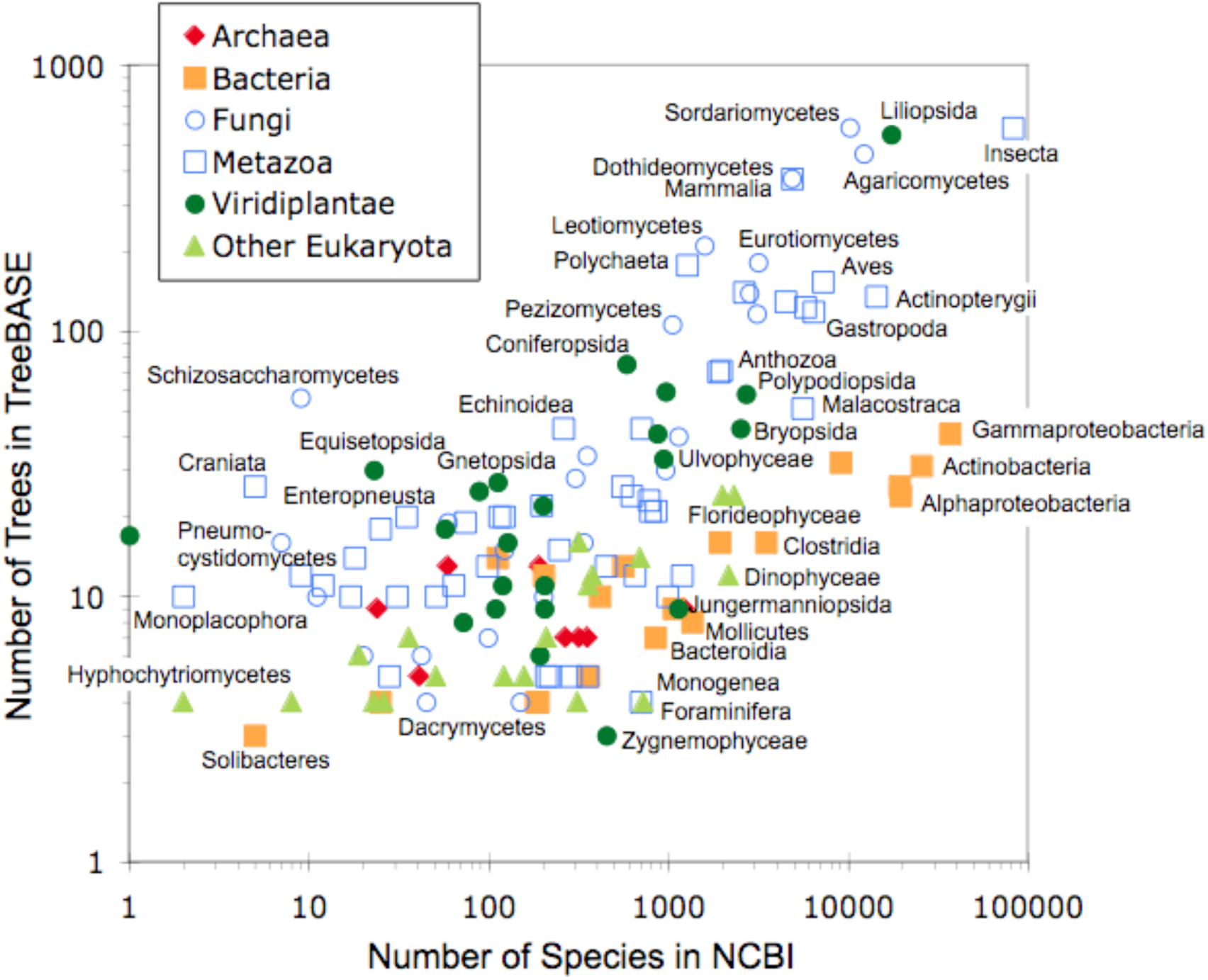
Biases in the taxonomic coverage of trees in TreeBASE. The abscissa is the number of trees in TreeBASE that include at least three taxa that belong to a given class; the ordinate is the number of species in NCBI belonging to this class. Although there is a correlation between the distribution of species in the NCBI taxonomy and the number of trees in TreeBASE, some classes have relatively more trees than others. Coordinates above the diagonal are classes that are better represented in TreeBASE than coordinates below the diagonal.

In addition to a web browser interface, TreeBASE offers two programmatic interfaces: the Open Archives Initiative Protocol for Metadata Harvesting (OAI-PMH) and the Phyloinformatics Web Services API (PhyloWS). The former service can be used to retrieve basic metadata for the set of studies created or modified after a certain date, for example:

http://treebase.org/treebase-web/top/oai?verb=ListRecords&metadataPrefix=oai_dc&from=2017-11-04T00:00:00Z

PhyloWS is a proposed mechanism for stateless phyloinformatic web services that largely conforms to the principles of Representational State Transfer (Fielding, R.T. and Taylor, R.N. 2000, Wilkinson, M.D. 2012) and that responds to queries written in Contextual Query Language (CQL). By default, responses are returned in RDF Site Summary (RSS 1.0) format, and therefore can be consumed and interpreted by any agent for automated phyloinformatic analysis that consumes RDF, including (but not limited to) news aggregator programs. For example, a user could set his or her news aggregator program to keep track of all new submissions to TreeBASE for papers published in the journal *Systematic Biology*:

http://purl.org/phylo/treebase/phylows/study/find?query=prism.publicationName=="Systematic+Biology"

TreeBASE uses globally unique Uniform Resource Identifiers (URIs) to reference studies, matrices, trees, and taxa. When URIs are dereferenced, useful metadata is returned in Resource Description Framework (RDF, http://www.w3.org/RDF/) so as to comply with Linked Data standards (Heath, T. and Bizer, C. 2011) in support of global semantic web integration. The following is an example URI to a study:

http://purl.org/phylo/treebase/phylows/study/TB2:S1925

Other formats are available, depending on the object. For example, study, matrix, and tree objects can be returned as NeXML or NEXUS by appending "?format=nexml" or "?format=nexus", respectively, to the end of the URI. Authors can use these links on their lab websites to point browsers directly to their data.

The main drawbacks of programmatic interfaces that operate on remote databases is that they tend to suffer from high latency and they tend to limit the user’s ability to customize queries. By offering TreeBASEdmp, a download of TreeBASE’s online holdings in a consolidated, simplified, and structured database format, we hope to improve the power and efficiency with which biologists can perform phyloinformatic analyses.

## INSTANTIATING THE DATABASE

The data to build TreeBASEdmp can be downloaded from https://figshare.com/projects/TreeBASEdb/37631 and decompressed into a file that is currently about 1.6 gigabytes in size called *treebasedmp.sql*. This SQL dump is designed for ingest into a PostgreSQL database. After installing PostgreSQL, a database can be created using the default template (i.e. with the command “CREATE DATABASE treebasedmp WITH TEMPLATE = template1;”) and the tables, sequences, constraints, data, and indices can then be ingested by executing the SQL dump (i.e. “\i treebasedmp.sql”). The resulting database takes up about 9.5GB of disk space. Since TreeBASE almost exclusively evolves by having data added but seldom deleted, the versioning of TreeBASEdmp is best identified by citing the DOI of the dataset. The last modification date of the database can be obtained with the following query: “SELECT max(lastmodifieddate) FROM study;”. We recommend that publications that make use of TreeBASEdmp in phyloinformatic research provide this paper’s citation as well as the DOI of the dataset.

## Database Model Walkthrough

The physical model for TreeBASEdmp is a simplified and consolidated version of the production database (Fig. 2), featuring only 14 relations as compared with the 99 relations in the production version.

**Figure 2.**
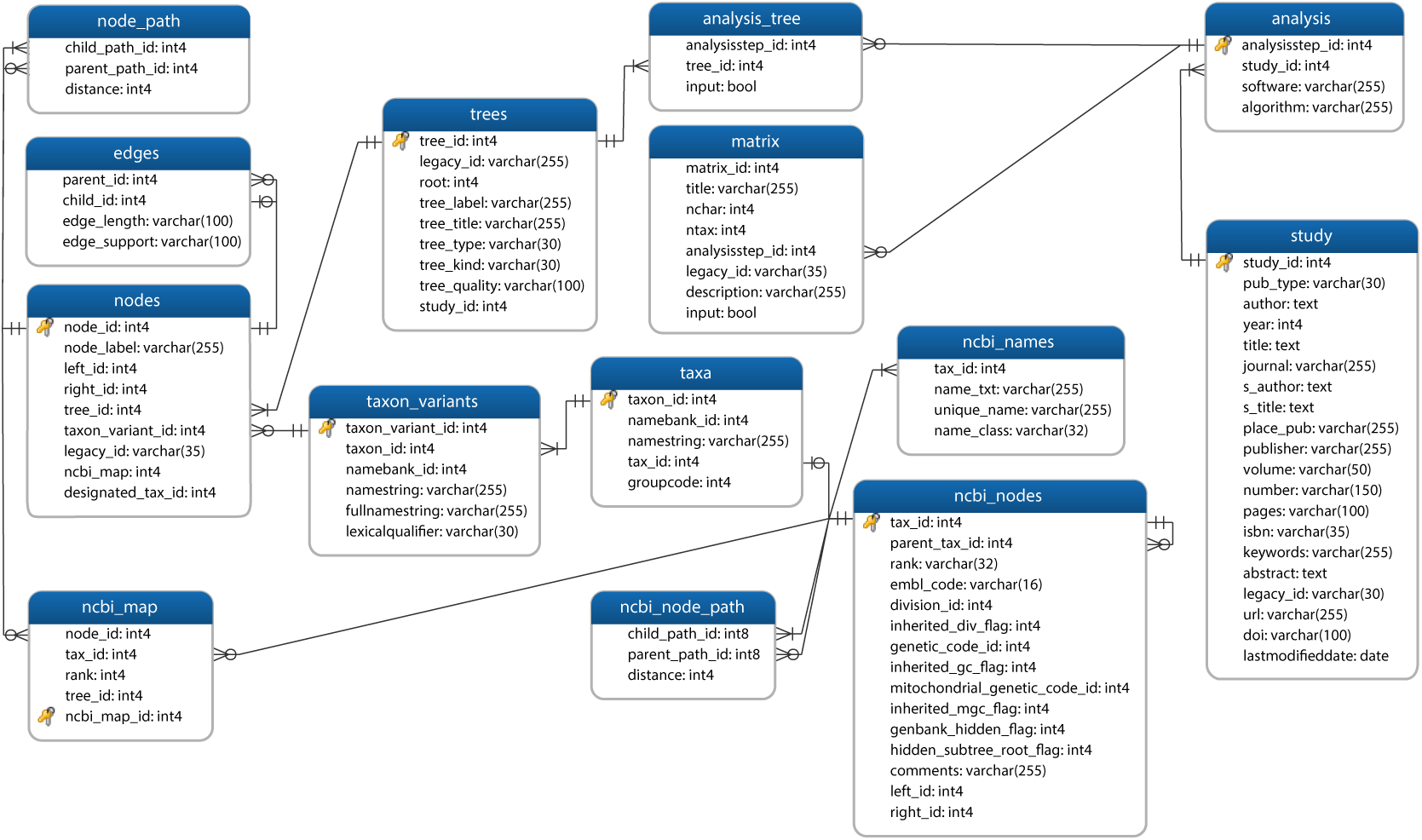
Entity Relationship Diagram describing the Physical Model of TreeBASEdmp using Crow Foot Notation.

Each publication archived in TreeBASE is represented by a record in the *study* relation and is assigned a unique integer identifier, *study.study_id*. The *study.pub_type* has one of three possible codes: “A” for a journal article; “I” for a book chapter, conference proceedings, or thesis chapter; and “B” for a book or thesis. If *study.pub_type* is “A,” the authors, year, article title, journal name, journal volume, journal issue number, article pages, abstract, and Digital Object Identifier are stored in *study.author, study.year, study.title, study.journal, study.volume, study.number, study.pages, study.abstract*, and *study.doi* respectively. If *study.pub_type* is either “I” or “B,” there is usually additional information in *study.place_pub, study.publisher*, and *study.isbn*. If *study.pub_type* is “I,” the book editors and book title are stored in *study.s_author* and *study.s_title* respectively. In addition, publication metadata can also be found in *study.keywords* and *study.abstract*. The column *study.legacy_id* is the unique identifier assigned to TreeBASE submissions prior to its migration to PostgreSQL in 2010.

Each *study* has one or more related *analysis* records, including metadata for software (*study.software*, e.g. “MrBayes”) and algorithm (*study.algorithm*, e.g. “neighbor joining”). Each *analysis* record has zero or more *matrix* and *trees* records. For most analyses, the *matrix* records are considered the inputs to the analysis (i.e. *matrix.input* = TRUE) while the *trees* records are considered the outputs (i.e. *analysis_tree.input* = FALSE). The *analysis* and *trees* are in a many-to-many relationship, joined by the many relation *analysis_tree*, thus allowing a given *tree* to belong to several *analysis* records. Analysis metadata includes *analysis.algorithm* (e.g. “parsimony,” “likelihood,” “bayesian inference,” etc.) and *analysis.software* (e.g. “PAUP,” “MrBayes,” “RAxML,” “MEGA,” etc.). While the production database also has many-to-many relations between *analysis* and *matrix*, here we have simplified this to a one-to-many relation, duplicating *matrix* entries as needed. The matrix relation does not actually store the alignment or data matrix, but only stores metadata about it, such as the dimensions, title, and description (i.e. *matrix.ntax, matrix.nchar, matrix.title, matrix.description*). Users who need access to the matrix data need to pull it from production – e.g. if the *matrix.matrix_id* is 553, resolving the following URL will retrieve the data in NeXML format: http://purl.org/phylo/treebase/phylows/matrix/TB2:M553?format=nexml.

The *trees* relation stores metadata about phylogenetic trees, including *trees.tree_label* (e.g. “Fig. 2”), *trees.tree_title* (e.g. “Phylogeny of Primates”), *trees.tree_type* (one of "Single," "Consensus," or "SuperTree"), *trees.tree_kind* (one of "Barcode Tree," "Language Tree," "Species Tree," or "Gene Tree"), and *trees.tree_quality* (e.g. one of "Alternative Tree," "Unrated," "Preferred Tree," or "Suboptimal Tree"). Each *trees* record has one or more *nodes*, and the *nodes.node_id* that serves as the root node is stored in *trees.root*. Even unrooted trees submitted to TreeBASE are stored with an arbitrarily designated root that corresponds to the outermost parentheses in the Newick notation of the submitted tree.

The *nodes* records are connected together in the tree network using an adjacency list table called *edges*, in which a node can serve as child (*edges.child_id*) to zero or one parent nodes, or as a parent (*edges.parent_id*) to one or more child nodes. The *edges* stores metadata associated with the child relation: the *edges.edge_length* is the branch length between the child node and the parent node; the *edges.edge_support* is the bootstrap, posterior probability, decay index, or any other support parameter associated with the child node.

The *nodes* are indexed to facilitate topological querying using both nested sets (Mackey, A. 2002) and transitive closure (Nakhleh, L., Miranker, D., et al. 2003, Trißl, S. and Leser, U. 2005). The nested set indices are integers incremented such that for a given tree, all nodes nested within a clade have a *nodes.left_id* that is equal or greater than the *nodes.left_id* of the clade node and at the same time less than the *nodes.right_id* of the clade node. Transitive closure uses the precomputed closure table *node_path* to store all possible paths between ancestral nodes (*node_path.parent_path_id*) and descendant nodes (*node_path.child_path_id*) in a given tree. The number of *edges* records needed to traverse a given path is stored in *node_path.distance*.

Each leaf node (identified by all *nodes* where *nodes.right_id* - *nodes.left_id* = 1) has text in *nodes.node_label* that represents the operational taxonomic unit (OTU) label as originally submitted to TreeBASE. This label is usually interpreted as a taxon and matched with a *taxon_variants* record using the taxon matching services of uBio. Beware that these matches may be incorrect (e.g. in the case of homonyms) or incomplete (e.g. the failure to match a *nodes.node_label* with a taxon). Each *taxa* record has one or more *taxon_variants*. The *taxon_variants.namestring, taxon_variants.fullnamestring*, and *taxon_variants.lexicalqualifier* are extracted from uBio, though in many cases *taxon_variants.namestring* reflects the label that a submitting author used in her or his data. Examples of *taxon_variants.lexicalqualifier* includes terms such as “chresonym,” “canonical misspelling,” and “anamorph.” The most canonical form of a taxonomic name – usually a binomial or trinomial – is expressed in *taxa.namestring*. If a *taxa* record appears to match a taxon in NCBI or uBio, it will show the respective identifiers in *taxa.tax_id* and *taxa.namebank_id*.

In most cases, the *taxa* relation has a *taxa.tax_id* that matches a record in NCBI taxonomy database, replicated here in *ncbi_nodes* and *ncbi_names.* These data are acquired from the NCBI (ftp://ftp.ncbi.nih.gov/pub/taxonomy/taxdmp.zip) at the time that the TreeBASE dump is created. As with the trees in TreeBASE, the NCBI classification tree is indexed using adjacency lists (*ncbi_nodes.tax_id* as child and *ncbi_nodes.parent_tax_id* as parent), nested sets (*ncbi_nodes.left_id* and *ncbi_nodes.right_id*), and transitive closure (*ncbi_node_path.child_path_id* and *ncbi_node_path.parent_path_id*).

There are three paths that can connect nodes in TreeBASE *trees* with taxonomic records in the NCBI taxonomy tables. In the first path, the leaf *nodes* (i.e. just the OTUs) can join with *taxon_variants* using *taxon_variants_id*, which can join with *taxa* using *taxon_id*, and finally join with *ncbi_nodes* using *taxa.tax_id*. In the second path, any record in *nodes* can have a many-to-many relation with *ncbi_nodes* using the *ncbi_map* joiner. While the former path is used in exact topological querying, the latter is used in generic topological querying, as later explained. The third path uses *nodes.designated_tax_id* to store species-level NCBI taxonomy identifiers, primarily to assist in supertree synthesis.

Since each OTU that maps to a taxon in *ncbi_nodes* can do so to a taxon of any rank, here we recalculate this relationship exclusively to the species rank. For OTUs that match to a subspecies, variant, cultivar, or any other rank below the species level, we store the parental species identifier in *nodes.designated_tax_id*. For OTUs that match to a genus, family, superfamily, or any other rank above the species level, we select the identifier of the descendant species that is most commonly found in TreeBASE trees. For example, an OTU with the taxon Primates will probably map to *Homo sapiens* because *Homo sapiens* is the most common primate in TreeBASE. Since most supertree methods perform best with maximal overlap of OTUs, this normalization to the common rank of species helps improve supertree synthesis. An alternative approach would be to substitute each higher taxon node with large polytomy of all descendant species, but this is more easily proliferated on the fly using a supertree assembly script.

## Meta-analysis Querying

TreeBASEdmp can be queried to investigate meta-analysis patterns in the field of phylogenetics, such as trends in usage of software programs, algorithms, taxonomic coverage, etc., or the phylogenetic output of any particular scientist. As a basic example, the following query (1) returns all of Michael J. Donoghue’s publications that included a morphological dataset that was analyzed using parsimony:

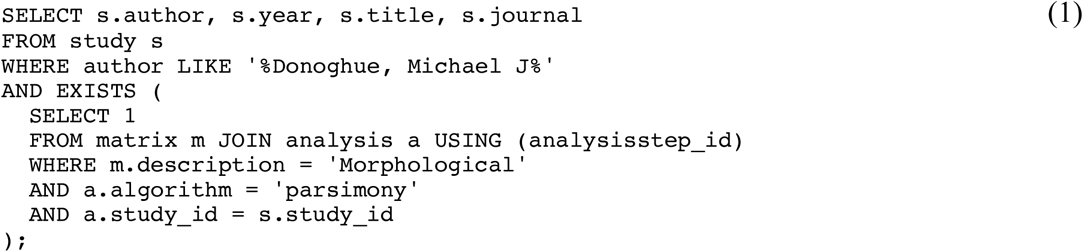

For a more complicated query, (2) illustrates how one can calculate the percentage of trees that result from using the program RAxML and the average size of trees, each on a year-by-year basis. This query uses four joined subqueries in the FROM clause: the first generates a list of years from 2006 to 2017; the second counts the number of trees in a given year; the third counts the number of OTUs (or leaf nodes) in a given year; and the last counts the number of trees per year that resulted from an analysis that uses the program RAxML.

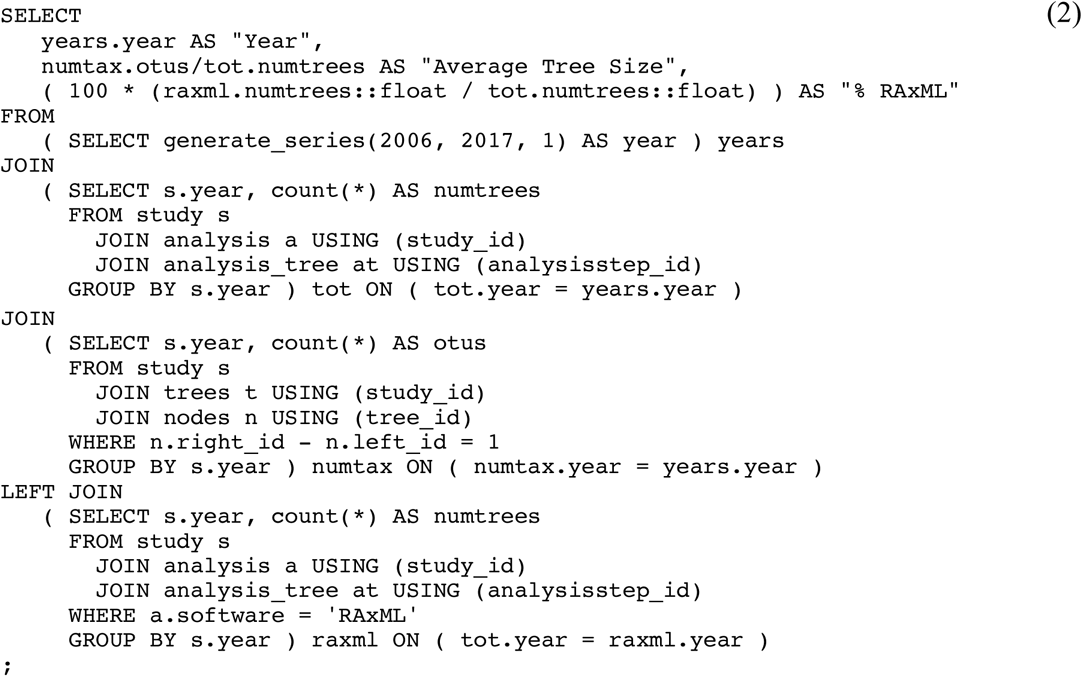

The results of this query (Fig. 3) show that the average size of trees in TreeBASE has been steadily rising from about 50 OTUs in 2006 to over 100 in 2017, and the percent usage of RAxML has also been rising. If the user wanted to demonstrate that RAxML is in some way responsible for larger analyses, the size of trees (along with, perhaps, the length of the data matrix, *matrix.nchar*) could be calculated for each type of software program. This is but one example of the kinds of meta-analysis queries that researchers may wish to perform. Note, however, that the annotation of metadata is largely a voluntary effort on the part of the submitter. For example, while submitters have the opportunity of designating the type of tree (e.g. single vs. consensus), or the kind of tree (e.g. species tree vs. gene tree), only a fraction of submitters take the effort to do this. Metadata that depends on voluntary effort is probably less reliable than computed metadata such as tree size, alignment length, or tree shape.

**Figure 3.**
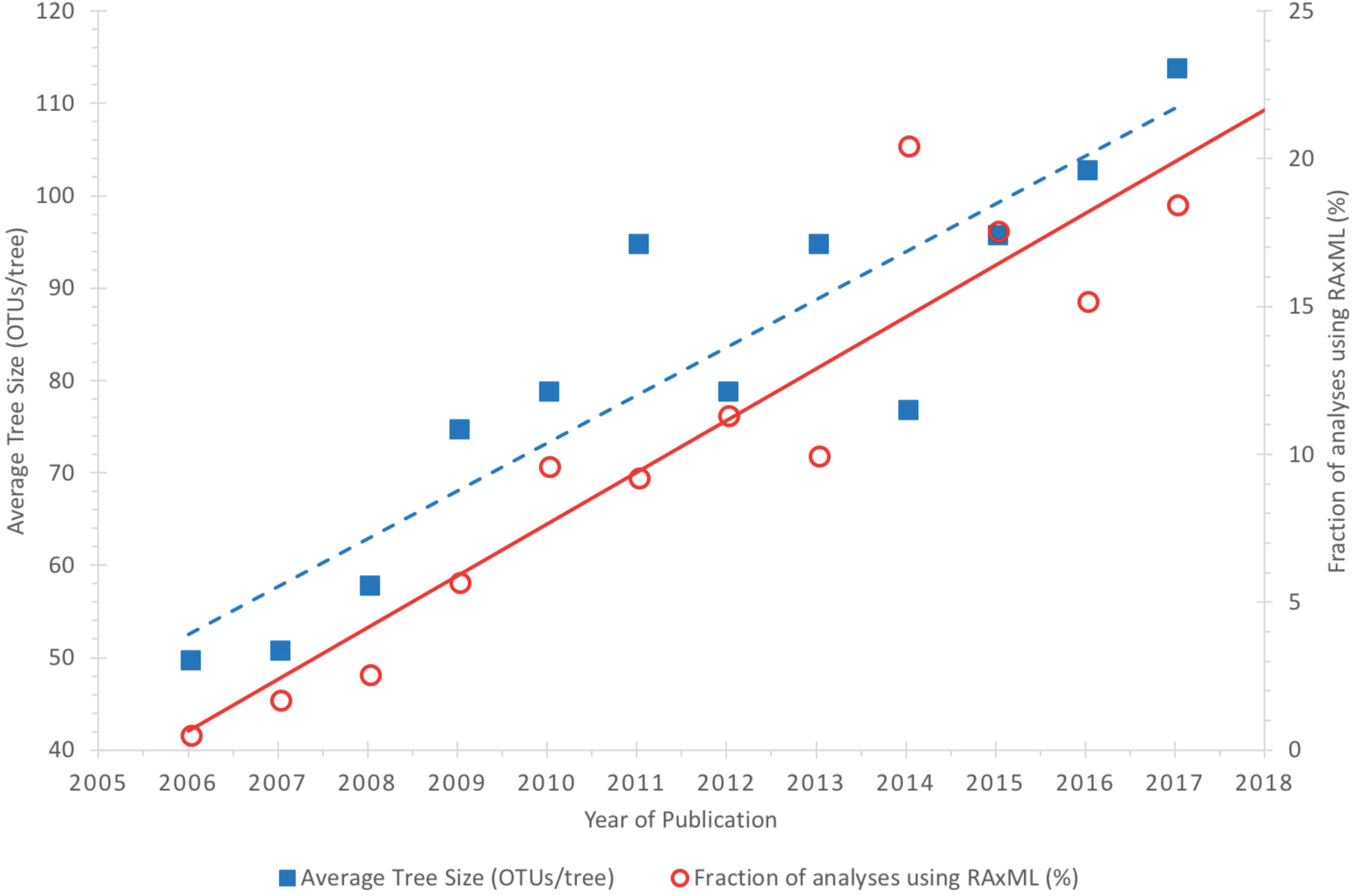
Results from our example meta-analysis query (2) indicate that both the average size of each tree and the fraction of analyses that use RAxML are rising steadily. Currently in TreeBASE the average tree size is about 110 OTUs and around 20% of analyses are performed using RAxML.

## Exact Topological Querying

An essential part of phyloinformatic research is in finding patterns of phylogenetic relationship. Whether for a large collection of trees or for a single very large tree, inspecting the trees visually is often impractical. The trees in TreeBASEdmp have been indexed with both nested sets and transitive closure to allow users to find trees or parts of trees that match a particular pattern of relationship.

### Queries using Nested Sets

In TreeBASEdmp, both the node records of TreeBASE trees and the NCBI taxonomy are indexed with *left_id* and *right_id* integers to facilitate nested set querying. The rationale behind using these indices is illustrated in Figure 4. As a simple example, we can start by querying the NCBI taxonomy tree to find all species nodes that descend from the node labeled “Aves” (i.e. all species of birds).

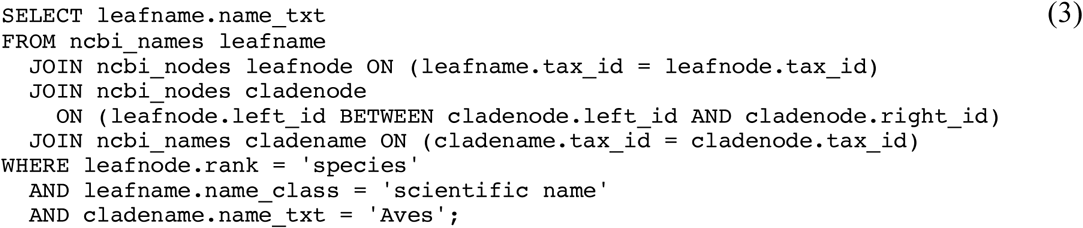

**Figure 4.**
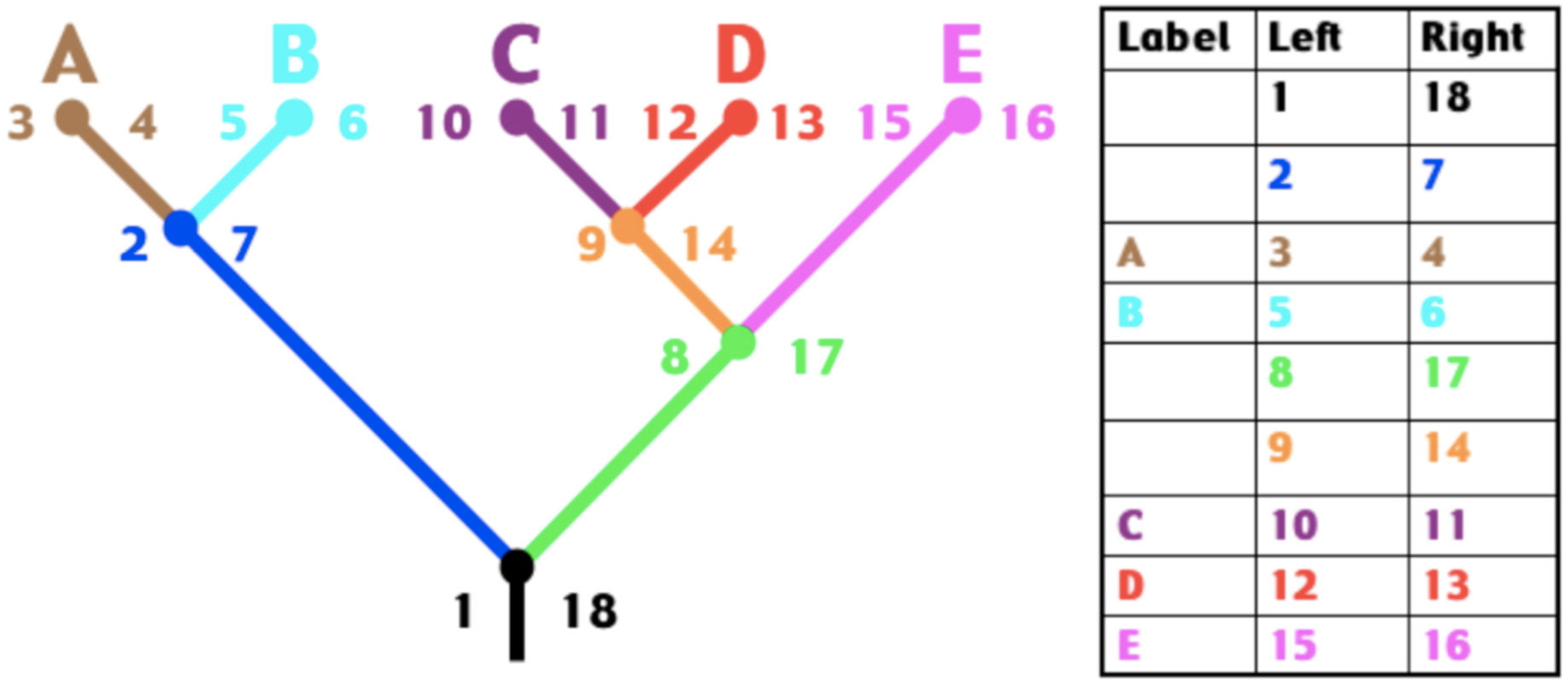
Illustration of nested set indexing. Each node on the tree on the left is labeled with *left_id* and *right_id* integers with values that are incremented in a depth-first traversal. Each row of the table on the right represents a node in the tree. Nodes that descend from a clade node have either their *left_id* or *right_id* integers greater than the *left_id* but less than the *right_id* of the clade node. Likewise, the ancestor nodes of a clade node have a *left_id* that is less than the *left_id* of the clade node and a *right_id* that is greater than the *right_id* of the clade node.

Query (3) creates a set of *ncbi_nodes* with the alias “leafnode” that have a *left_id* value between the *left_id* and *right_id* of a node (here with the alias “cladenode”) that has a *tax_id* that matches the taxon Aves. The result indicates that there are just shy of 10,000 species of birds in the NCBI taxonomy. We can modify (3) into a series of subqueries to find all trees in TreeBASE that have at least one bird taxon in the tree but have neither a mammal nor a crocodile taxon in the tree, as follows:

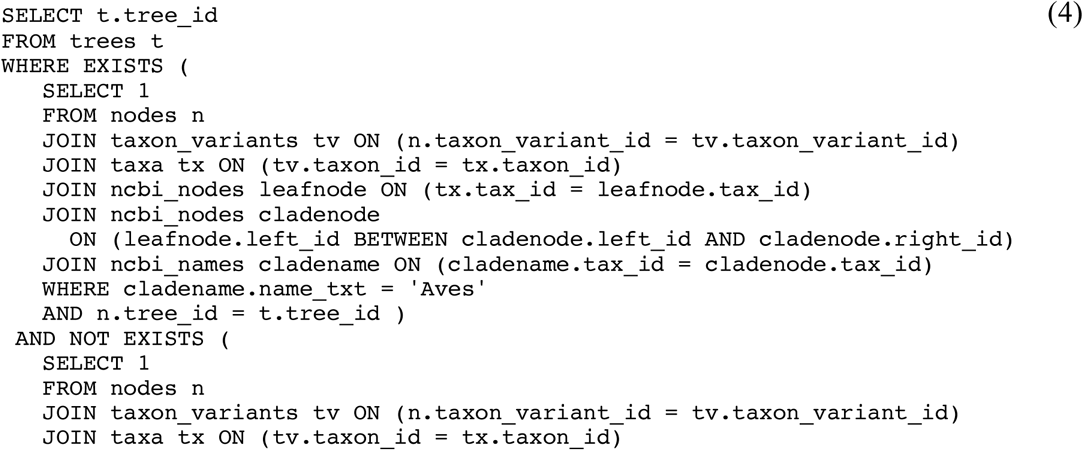

Executing query (4) indicates that over 360 trees match these criteria.

A key step in finding topological patterns in a single tree or a collection of trees is to identify the most recent common ancestor (MRCA) for a set of leaf nodes. Nested set indexing can operate on a single tree or a collection of trees to identify the MRCAs given a set of leaf nodes. The following query identifies the MRCA nodes in each tree that has leaf labels with both *Hippopotamus amphibius* and *Sus scrofa*:

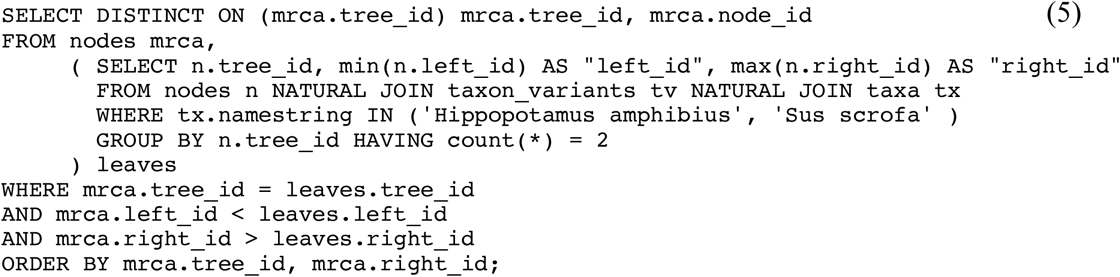

The subquery *leaves* finds all nodes in trees that have all the listed taxon names (and if there were more than two in the list, the integer in the GROUP BY aggregate function should be adjusted accordingly). Using this pair of nodes in each tree, the query then finds all common ancestors, and from these it picks the most recent one by ordering them by *right_id*. Note that the use of DISTINCT ON is peculiar to PostgreSQL.

We can reuse query (5) as a subquery to find any trees where *Sus scrofa* is nested within the MRCA of *Hippopotamus amphibius* and *Tursiops truncatus*. If a tree is found, it would contradict the currently accepted notion that hippos and cetaceans are sister groups to the exclusion of other extant artiodactyls, such as pigs.

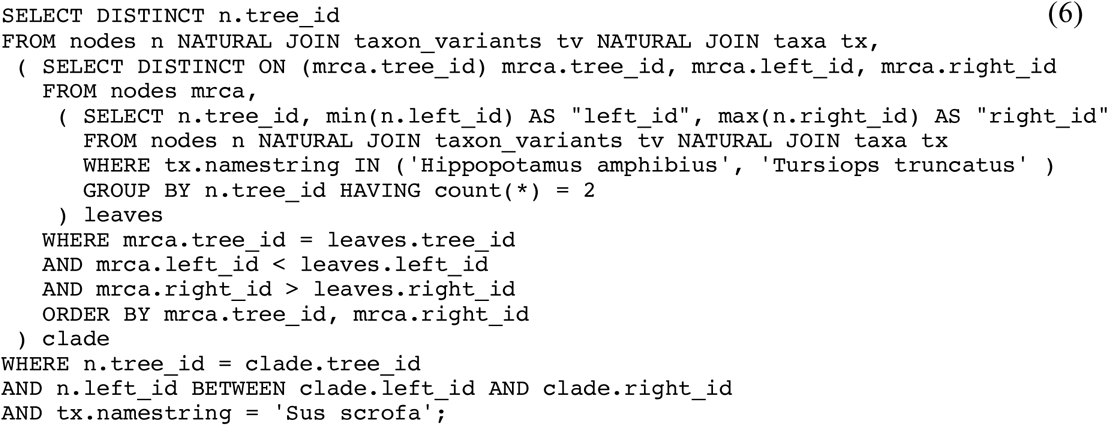

In query (6), the subquery with the alias *clade* lists all the MRCAs of hippos and dolphins. It’s then a matter of finding whether the *left_id* of any nodes labeled with *Sus scrofa* falls between *clade.left_id* and *clade.right_id*. With some 20,000 trees in the database, query (6) retrieves this topological pattern in about 0.3 seconds running on a typical consumer notebook computer.

### Queries using Transitive Closure

Precomputing a closure table of all possible ancestor-descendant paths proliferates a lot of records (in the case of TreeBASEdmp, about a quarter of a billion records) but further expands our ability to design queries for finding topological patterns. The rationale for this approach is illustrated in Figure 5, where the closure table (Fig. 5B) includes the same information as in an *edges* table, but goes beyond just parent-child pairs to also include parent-grandchild, parent-great-grandchild, and so forth. The common ancestors between any two leaf nodes can be found by querying for these leaf nodes while requiring matching *parent_id* values in the closure table. In TreeBASEdmp, closure tables are computed for both the TreeBASE trees and the NCBI taxonomy tree. In the following example, we query the NCBI taxonomy tree to get a list of all the common ancestors to *Pan troglodyte*s and *Bos taurus*:

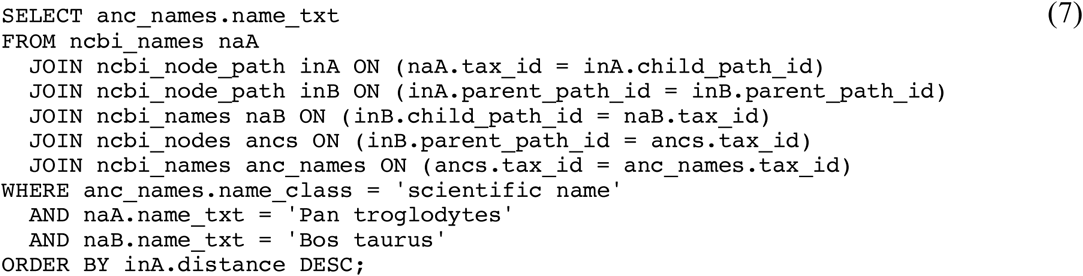

**Figure 5.**
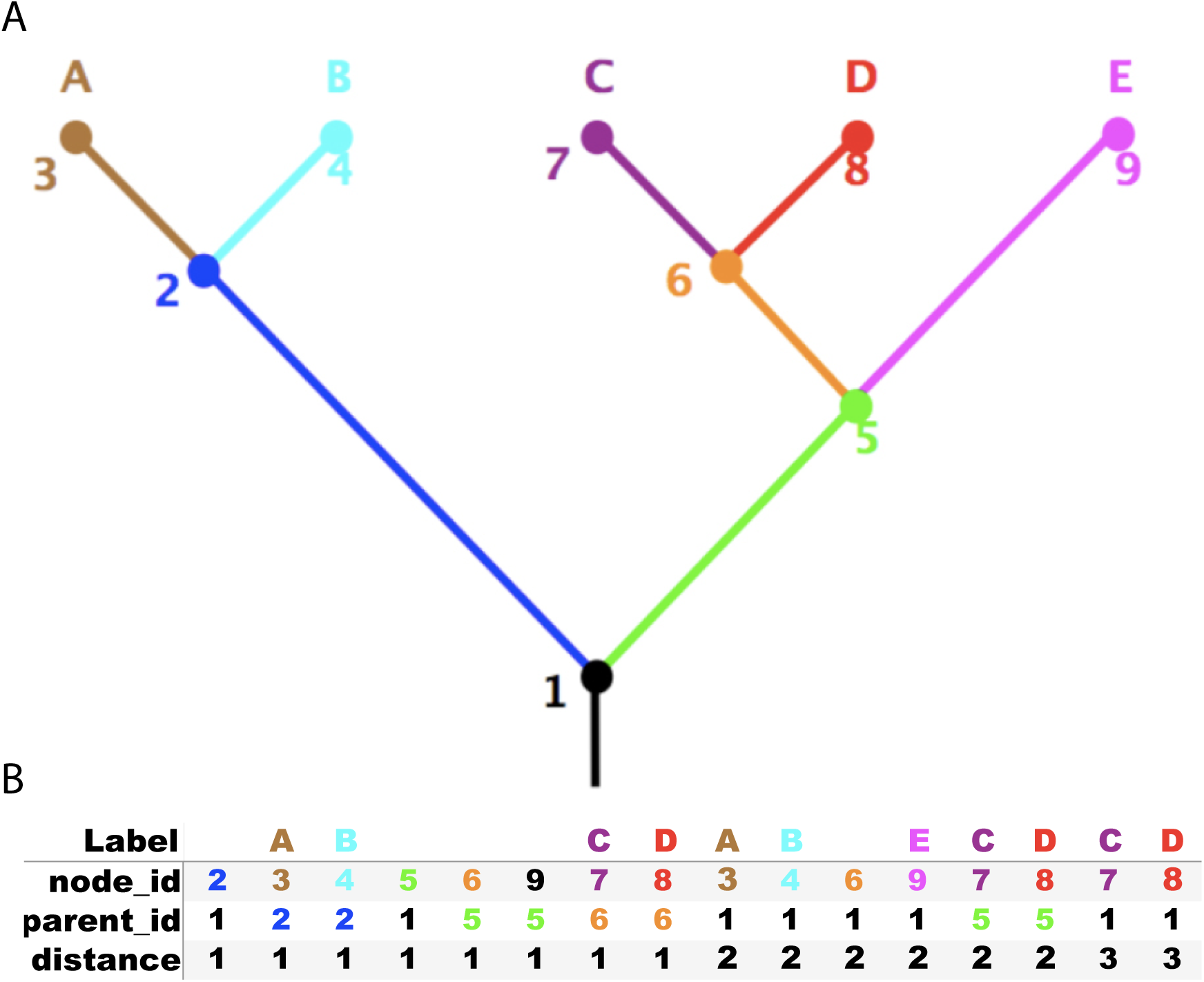
Illustration of transitive closure indexing: an example tree (A) and the corresponding closure table (B). The closure table lists all possible ancestor-descendant node paths, with the number of edges indicated as the *distance*. In this example, the first eight records are the same as what would be stored in the *edges* table, i.e. parent-child records. The remaining eight records represent longer paths, i.e. grandparent-child records, great grandparent-child records, etc.

The result is a list of higher taxa in order starting from the root (i.e. life, cellular organisms, Eukaryota, Opisthokonta, Metazoa, etc.), and therefore the first name is the oldest common ancestor and the last name is the MRCA. We can reuse query (7) as a subquery to produce a more interesting result, such as find the oldest common ancestor of chimpanzee and cow that is not a common ancestor of a frog and salamander:

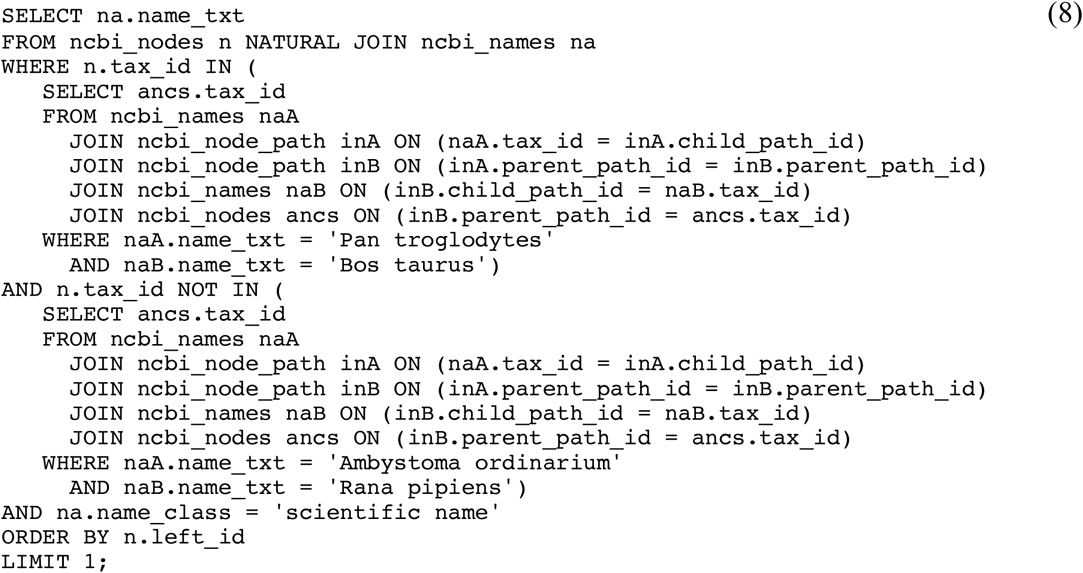

Naturally, the result of query (8) is “Amniota” – a fairly trivial calculation when applied to this classification tree, but an essential query when applied to phylogenetic trees in support of generic topological querying, as we will see later on.

Using the closure table *node_path*, we can rewrite query (6) to perform the same function, but here using the closure table instead of nested set indexing, i.e. query (9) returns all trees where pig is nested within the MRCA of hippo and dolphin:

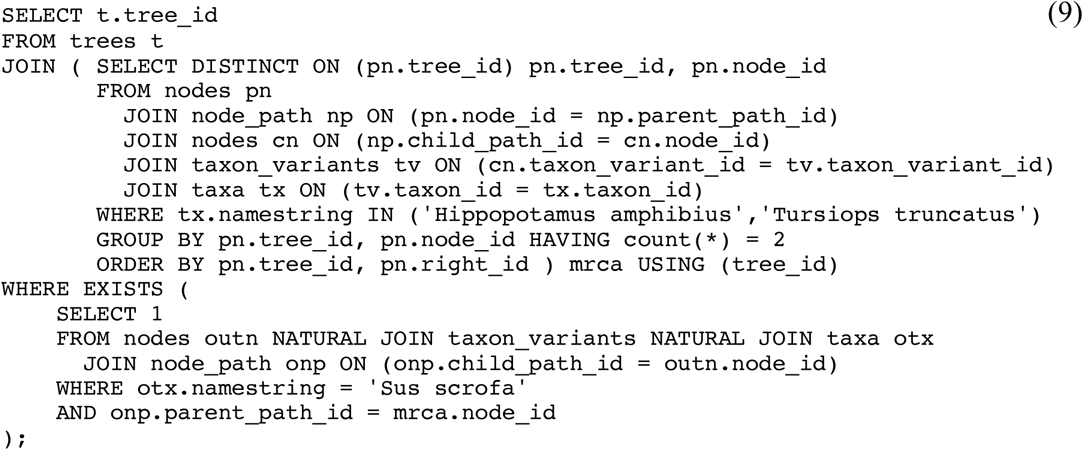

The subquery in the FROM clause returns the MRCA nodes for hippo and dolphin, while the subquery in the WHERE clause limits the result set to just those trees that also have *Sus scrofa* as a descendant of the MRCA nodes.

## Generic Topological Querying

With “exact” topological querying, the taxonomic labels identifying leaf nodes are exactly specified. This approach is most effective when searching a set of trees where the same set of leaf node labels are used throughout. For example, if you wanted to filter a set of trees that resulted from a Bayesian MCMC run to estimate the posterior probability of a particular topological pattern, you would count on the fact that each tree shares the same set of taxon labels. However phylogenetic knowledge on the whole is expressed as a large heterogenous collection of trees varying in size and depth, where few if any share the same set of taxonomic labels. There are two main challenges here: one is with semantic heterogeneity in taxonomic identifiers, and the other is in querying a particular phylogenetic hypothesis broadly conceived and regardless of which taxa are used to represent the leaf nodes.

The problem of semantic heterogeneity takes many forms: different ways of writing the same taxonomic name (i.e. misspellings, vernacular variants, and lexical variants), names that may or may not refer to the same taxon (i.e. heterotypic and homotypic synonyms), identical names that refer to different taxa (i.e. homonyms), and differences in taxonomic application (i.e. differences in circumscription). Various solutions to resolving semantic heterogeneity have emerged (Page, R.D.M. 2008, Boyle, B., Hopkins, N., et al. 2013, Rees, J.A. and Cranston, K. 2017), but thus far there are no perfect solutions, especially for subjective synonyms and differences in taxonomic application. TreeBASE attempts to resolve semantic heterogeneity problems using uBio name services together with synonym resolution data in the NCBI classification. The many-to-one relationship between *taxon_variants* and *taxa* relations expresses the notion that there are many ways that the same taxon can be expressed in the trees submitted to the database. Furthermore, the inclusion of junior synonyms in the *ncbi_names* relation can be used to help resolve objective synonyms.

However, even if the meaning of taxonomic identifiers can be resolved accurately, there still remains the problem of expressing the same phylogenetic pattern despite using different taxa as OTUs. This problem is illustrated in Figure 6, where the OTU labels of trees A and B are different even though both trees express the same general phylogenetic hypothesis. To support *generic topological querying*, where topology can be queried despite trees using different OTUs, we have computed the relation *ncbi_map* to store a hypothesized equivalence between higher taxa in NCBI and as many nodes as possible in the trees. The root node of each tree maps to an NCBI higher taxon representing the MRCA of all the taxa in the tree. In this example, the root nodes of both Fig. 6A and Fig. 6B map to the Laurasiatheria. For all remaining nodes in the tree, we apply query (8) in an attempt to map them to NCBI higher taxa representing common ancestors of the ingroup so long as none are parent to any non-ingroup taxa. For both Fig. 6A and Fig. 6B, all remaining nodes map to the same set of oldest common ancestors of their ingroups. Using these mappings, the trees in Figs. 6C and 6D are now easily searched for topological patterns. This approach is limited by the resolution of the taxonomic classification – e.g. since the NCBI classification has no higher taxon to represent the clade that unites hippos and cetaceans, such nodes cannot be mapped.

**Figure 6.**
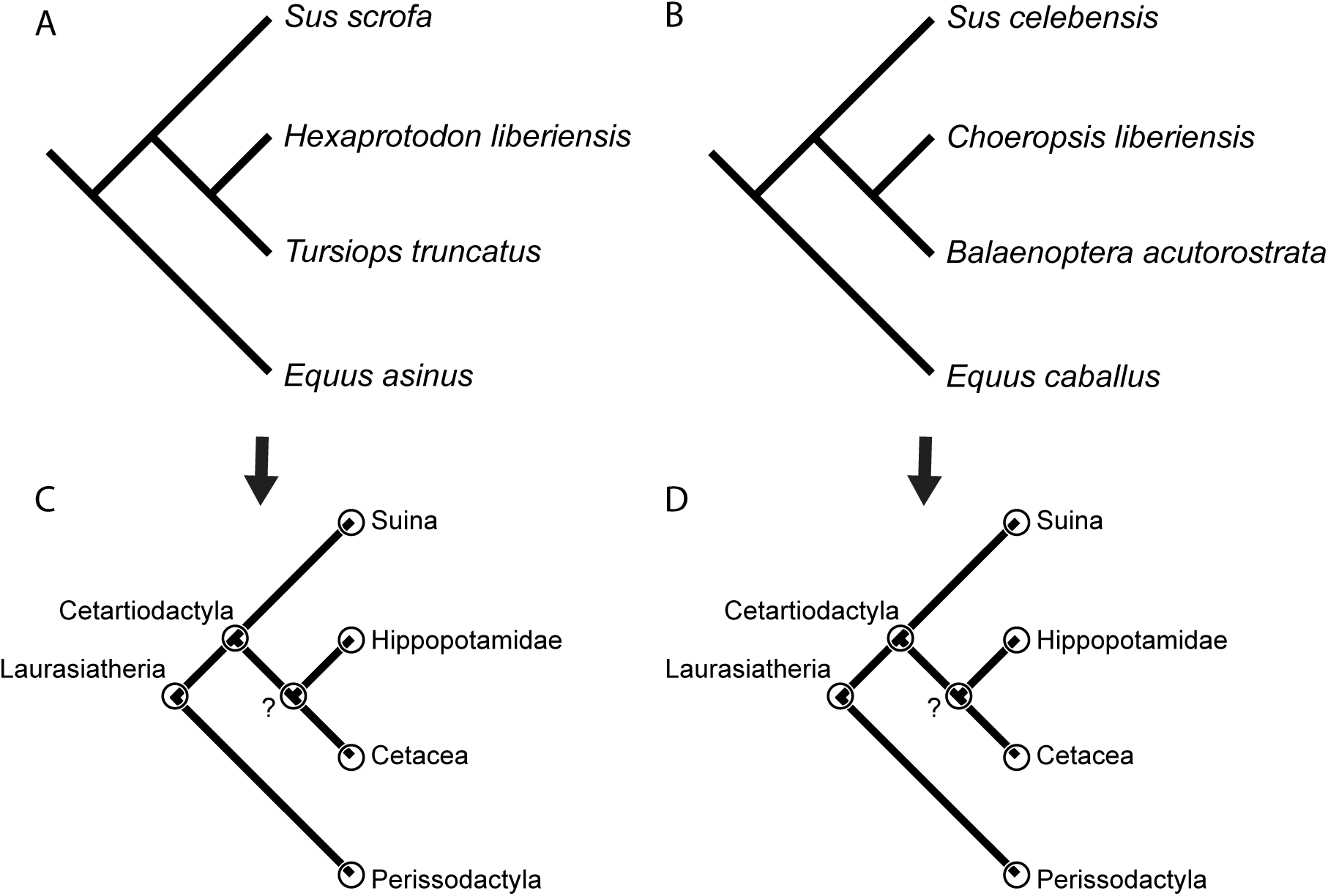
Illustration of the difficulty of performing topological querying on trees with leaf node labels that use either semantic heterogeneity or taxonomic heterogeneity. The trees figured in (A) and (B) are essentially stating the same phylogenetic hypothesis, yet none of their leaf labels match up. In the case of the pygmy hippo, both *Choeropsis liberiensis* and *Hexaprotodon liberiensis* are objective synonyms of the same taxon but use different names. For all remaining leaf nodes, trees A and B use different species as OTUs. To perform generic topological querying, each node, except for the root node, is mapped to a higher taxon name using query (8) – i.e. the oldest common ancestor of the ingroup to the exclusion of all other non-ingroups. The root node is mapped to the MRCA of all taxa in the tree. Performing this operation on trees A and B results in mapped trees C and D respectively. Once mapped in this way, it is clear that the trees were stating the same phylogenetic hypothesis, and both trees can be recovered using the same topological query.

Using the *ncbi_map* relation, we can now query for trees that match our phylogenetic question regardless of which specific species were used. Here we will search for trees that match the modern molecular hypothesis for artiodactyls – i.e. all trees having a clade of hippos and cetaceans but do not have any pigs, camels, or ruminants nested within this clade:

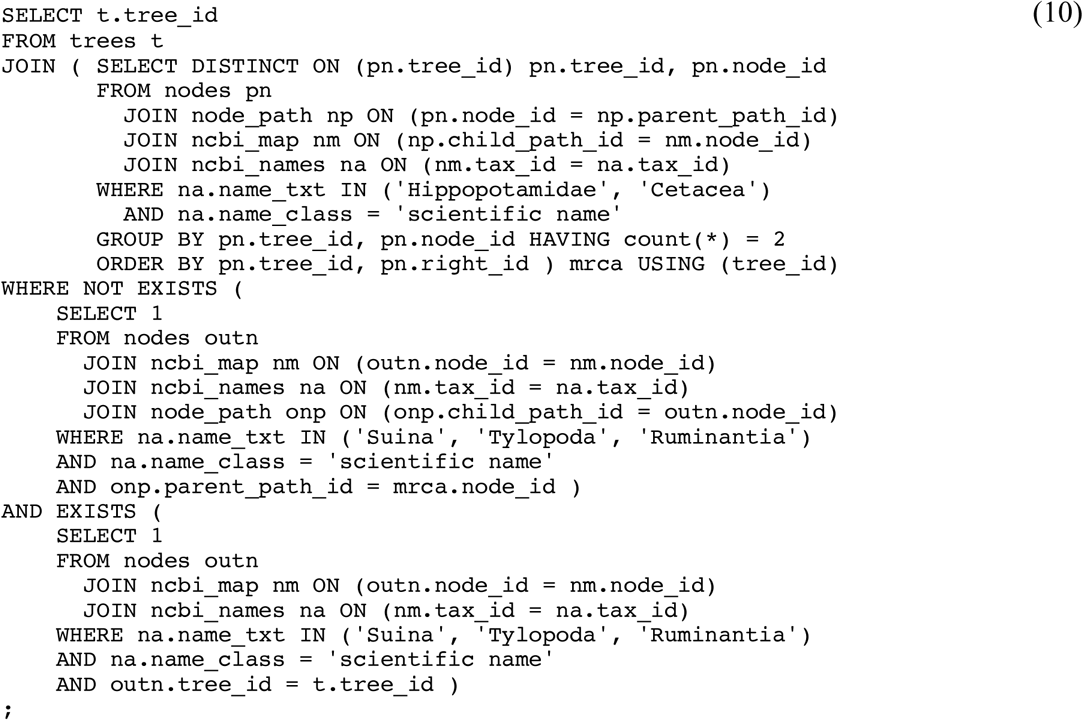

The subquery in the FROM clause returns all MRCA *node_id* integers for cetaceans and hippos. In the WHERE clause, the first subquery excludes any trees where a pig, camel, or ruminant is found to descend from any of these MRCA nodes. This query would not be very meaningful if it returned trees that only had cetaceans and hippos, so the second subquery is used to require that the trees have at least one pig, camel, or ruminant.

## Rendering Trees

Complex queries typically result in a set of trees or nodes that can then be used in further analyses, such as building supertrees or calculating tree-to-tree distance metrics. One option is to use the values in *trees.tree_id* as part of an URL to retrieve trees serialized in NEXUS or NeXML. In the following example, if the value of *trees.tree_id* is 1000, the NEXUS-formatted tree will be returned from this URL:

http://purl.org/phylo/treebase/phylows/tree/TB2:Tr1000?format=nexus

This solution is not ideal because of latency in download speeds and the limited options in how the trees are formatted. A better solution is to build trees directly from *TreeBASEdmp* using a procedural language with appropriate phylogenetic and database connectivity libraries. As an example, Appendix S1 is a Perl script that takes a list of *nodes.node_id* values as starting nodes and for each creates tree objects using the Bio::Phylo library. Using this library, various calculations can be made over the resulting collection of trees and different serializations can be generated. In our example we simply write the set of trees to a file in NEXUS format.

## Building Supertrees

A common way to build a synthesis of phylogenetic knowledge is to produce a supertree based on a set of input trees (Bininda-Emonds, O.R. 2004). Supertrees of species are useful for a variety of purposes – examples include using independent contrasts to factor out the effect of phylogeny in hypothesis testing, or for reconciling species trees with gene trees to infer gene duplication events. For the most part, supertree methods improve when there is a greater degree of OTU overlap among input trees. This requirement is addressed in TreeBASEdmp by normalizing OTUs to the species rank using *nodes.designated_tax_id*, as previously explained. However, users are not obliged to use *nodes.designated_tax_id*, and are free to normalize OTUs using other methods.

We offer a Perl script in Appendix S2 to show how supertrees can be calculated using matrix representation with parsimony (MRP) (Baum, B.R. 1992, Ragan, M.A. 1992). The input parameter is any higher taxon name (e.g. “Primates”). The output is a NEXUS file containing a matrix representation for subtrees starting at internal nodes that map to the higher taxon, as well as entire trees, each with a root node that descends from the higher taxon.

Example results are shown in supplemental figure Fig. S1. Tree (A) is a supertree of the mammal order Primates based on 96 trees resulting in a matrix of 319 species and 2041 characters, inferred using a “qnelsen” parsimony search with TNT (Goloboff, P.A., Farris, J.S., et al. 2008). Tree (B) is a supertree of the asterid order Dipsacales based on 132 trees resulting in a matrix of 406 species and 3319 characters. Tree (C) is a supertree of the lepidopteran superfamily Papilionoidea based on 172 trees resulting in a matrix of 758 species and 5026 characters. In the case of (B) and (C), the trees were inferred using a heuristic parsimony search with PAUP (Swofford, D.L. 2001) retaining a maximum of 10,000 trees. The resulting trees are each a majority rule consensus, with clade frequencies indicated as percentages.

Naturally, users are not limited to MRP. Minor modification of the script in Appendix S2 could output trees instead of characters for analysis using other methods, such as MinCut (Semple, C. and Steel, M. 2000, Page, R.D.M. 2002), QFit (Reaz, R., Bayzid, M.S., et al. 2014), etc. To the extent that TreeBASE data are incomplete for the purposes of supertree construction, we suggest that users submit any missing trees to TreeBASE via the web portal, and then download the next TreeBASEdmp for supertree assembly.

## Availability

The TreeBASEdmp is available for download from the Figshare project https://figshare.com/projects/TreeBASEdb/37631, where individual releases are identified by their creation date stamp in ISO-8601 format, e.g. 2018-08-22 for the release that was posted on August 22nd, 2018. To verify the integrity of the download, an MD5 checksum is also available. The data dump is made available under a CC0 license, i.e. it can be reused for any purpose without restriction. Note, however, that these permissions apply solely to the facts (i.e. scientific data and phylogenetic knowledge) contained in the dump, and not to the scholarly publications and associated artwork that discuss these facts.

## Funding

TreeBASE has received support from the National Science Foundation (grant numbers DEB 9318325, EF 0331654, IIS 0629702, IIS 0629846, and DBI 0743720). TreeBASEdmp development was supported by Yale-NUS (grants SUG and IG15-SI101).

## Acknowledgements

TreeBASE oversight is provided by the Phyloinformatics Research Foundation Inc. and has received institutional support, in the form of bandwidth, hosting, and hardware, from Harvard University Herbaria, Leiden University, University at Buffalo, Yale Peabody Museum, the National Evolutionary Synthesis Center (NESCent). TreeBASE is currently hosted at the Naturalis Biodiversity Center, Leiden, The Netherlands.

